# Molecular Mechanism of Brassinosteroids Perception by the Plant Growth Receptor BRI1

**DOI:** 10.1101/794750

**Authors:** Faisal Aldukhi, Aniket Deb, Chuankai Zhao, Alexander S. Moffett, Diwakar Shukla

**Author notes:** These authors contributed equally to this work (Faisal Aldukhi, Aniket Deb).

## Abstract

Brassinosteroids (BRs) are essential phytohormones which bind to the plant receptor, BRI1, to regulate various physiological processes. The molecular mechanism of the perception of BRs by the ectodomain of BRI1 remains not fully understood. It also remains elusive why a substantial difference in biological activity exists between the BRs. In this work, we study the binding mechanisms of the two most bioactive BRs, brassinolide (BLD) and castasterone (CAT) using molecular dynamics simulations. We report free energy landscapes of the binding processes of both ligands as well as detailed ligand binding pathways. Our results suggest that CAT has lower binding affinity compared to BLD due to its inability to form hydrogen bonding interactions with a tyrosine residue in the island domain of BRI1. We uncover a conserved non-productive binding state for both BLD and CAT, which is more stable for CAT and may further contribute to the bioactivity difference. Finally, we validate past observations about the conformational restructuring and ordering of the island domain upon BLD binding. Overall, this study provides new insights into the fundamental mechanism of the perception of two most bioactive BRs, which may create new avenues for genetic and agrochemical control of their signaling cascade.

## Introduction

Plants, being immobile, face considerable challenges in adapting to their changing environments in order to grow and survive.^1^ Environmental signals, such as temperature and light, must then be sensed and acted upon by the plant intracellular system.^2^ This process of generating cellular responses to external environmental stimuli is facilitated via phytohormones. Brassinosteroids (BRs) represent an important category of phytohormones, which influence a wide range of physiological processes critical for plant growth and development.^3^ Mutants with defects in the biosynthesis or signaling of BRs show multiple inadequacies such as dwarfism, delay in flowering, reduced germination of seeds, low fertility levels and improper stomatal distribution.^4^ BR signaling starts with the Brassinosteroid Insensitive 1 (BRI1), a receptor kinase present at the cell surface. The extracellular domain of BRI1 recognizes BRs which leads to heteromerization with the co-receptor BRI1-Associated Receptor Kinase1 (BAK1), a member of the Somatic Embryogenesis Receptor Kinase (SERK) family of proteins. This is followed by transphosphorylation of the intracellular kinase domains of BRI1 and BAK1, which triggers a downstream signaling cascade eventually leading to the expression or suppression of important genes. ^1,4–7^

The perception of BRs by the BRI1 receptor is a crucial step in the BR signaling pathway.^8^ Previous studies have provided insights into the structure of this important plant receptor.^9–12^ BRI1 is localized in the plant cell membrane and consists of an extracellular ligand-binding domain, a single transmembrane helix and an intracellular kinase domain with serine/threonine specificity. The extracellular domain consists of 25 leucine-rich repeats (LRRs) and assumes a right-handed superhelical structure (Figure 1). ^9,11^ A notable feature of this part of the receptor is the presence of an island domain, which consists of 70 amino acid residues between the LRRs 21 and 22.^9,12^ The crystal structure of BRI1 with brassinolide (BLD), the most active bassinosteroid (Figure 1A), bound reveals that LRRs 21-25 along with the island domain form the binding pocket for BLD.^9,12^ It was suggested that BLD binding does not induce much conformational change in the LRR core of BRI1.^9,12^ However, the island domain becomes increasingly ordered upon the binding of BLD, allowing for the activation of the receptor.^9,12^ When BLD is bound to BRI1, the A-D rings of brassinolide are positioned in a hydrophobic groove between the LRR core and the island domain while its alkyl chain goes into a small pocket formed by LRRs 21, 22 and the island domain (Figure 1C). Interactions of BLD with the residues of the island domain and LRR core appear to be essential for BLD binding. Mutations in the residues of the island domain as well as of the nearby LRRs have been reported to adversely affect BLD binding and consequently the downstream signaling. ^8,9,12,13^ Thus, the island domain and the surrounding LRR core is indispensable for brassinolide perception.

**Figure 1:**
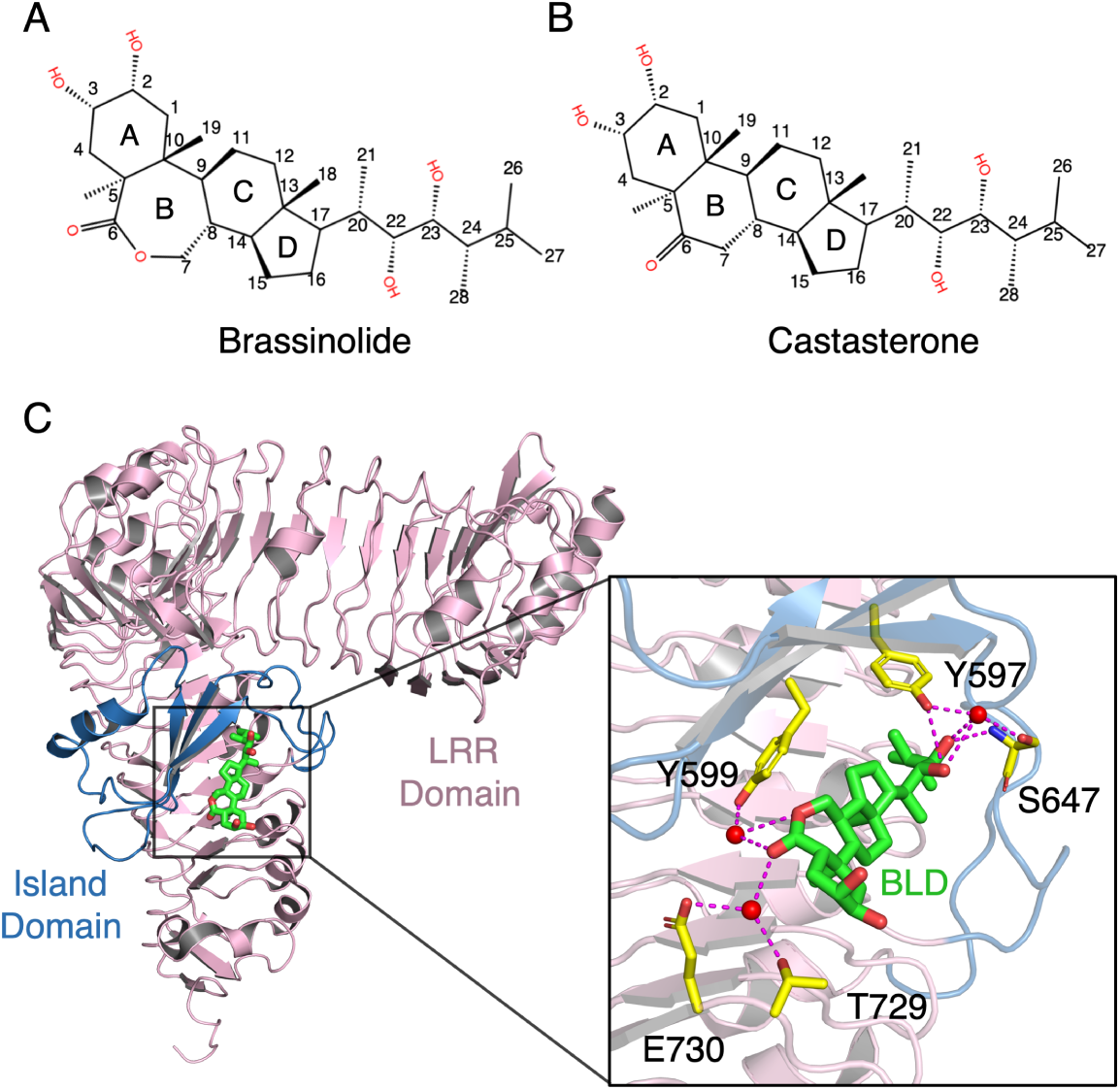
The chemical structures of (A) BLD and (B) CAT. The difference between the two molecules is the extra oxygen in the B ring of BLD. (C) Crystal structure of the BRI1-BLD complex (PDB ID: 3RGZ). The LRR core of the receptor is shown in pink and the island domain in sky blue. BLD is shown as green sticks while water molecules are shown as red spheres. Hydrogen bonds (magenta), either direct or water-mediated, between BLD and the residues of the island domain and LRR core of BRI1 are shown in the enlarged view.

Within the family of naturally synthesized BRs, BLD and castasterone (CAT, Figure 1B) are the most widely distributed in plants.^14^ In *Arabidopsis thaliana*, they have been isolated in the shoots, seeds and root calluses. BLD is isolated at higher quantities in the root callus while CAT is the dominant BR in the shoots.^15^ Previous results have classified BLD as the most bioactive BR.^16^ It is structurally a steroid with a unique lactone group in its B-ring and vicinal diols both in its A-ring and alkyl chain (Figure 1A).^17^ CAT is the second most potent BR and the immediate precursor of BLD in BR biosynthesis.^16,18^ It is structurally almost identical to BLD with the only difference being the absence of an oxygen atom between C6 and C7 carbons in the B-ring (Figure 1B).^19^ Although CAT can also act as a biologically active BR, it has been reported to be much less active than BLD, with difference ranging from 5-fold in rice lamina inclination test,^20,21^ 10-fold in the mung bean explant test^22^ to a 100-fold in the pinto-bean second internode test.^23^ It remains unclear how these two hormones with strikingly similar structures can have such a considerable difference in their biological activity.

Despite advances in mechanistic understanding of BR signaling, a large number of questions remain unanswered regarding the perception of BRs. The structural characterization of BRI1 via crystallographic methods has given crucial insights into the mode of action of BLD. However, a detailed atomic view and quantitaive thermodynamic and kinetic characterization of the binding of BLD to BRI1 is still lacking. Furthermore, there is limited information about the binding mechanisms of BRs other than BLD such as CAT. Particularly, the binding pose for CAT remains to be characterized. Biochemical assays have provided significant information about the relative differences in the biological activities of BRs. However, the molecular origin of the activity differences between BLD and CAT remains unclear. Therefore, a complete dynamic view and a detailed thermodynamic and kinetic investigation on the binding processes of BRs is required to fully understand their perception mechanism.

To answer these vital questions, one requires a tool which can describe the receptor conformations and interactions with the ligands at a high spatial-temporal resolution. Molecular dynamics (MD) simulations have gained considerable popularity as a tool for studying the dynamic behavior of biological systems at the atomic level. However, MD simulations alone typically require significant computational time to fully characterize long timescale biological processes, such as ligand binding, which occur in microseconds to milliseconds. One way to counter this is to combine MD simulations with adaptive sampling and Markov State Models (MSMs).^24–27^ This computational framework allows us to efficiently sample rare biological events and construct a discretized kinetic network model to describe the thermodynamics and kinetics of biological processes.^28–30^ This approach of MD simulations followed by MSM analysis has previously been successful in predicting protein-ligand binding processes^31,32^ and elucidating important conformational changes in proteins. ^33–36^

In this study, we employed the combined approach of MD simulations and MSM analysis to answer important questions regarding the perception of BLD and CAT by BRI1 in *Arabidopsis thaliana*. We have performed *∼*58 *µ*s and *∼*44 *µ*s MD simulations to capture the binding of BLD and CAT to BRI1, respectively. We report free energy landscapes to describe the thermodynamics of the binding of both ligands and identify the important binding intermediate states. We have replicated the bound pose for BLD including the important interactions with BRI1 residues as elaborated by the crystal structures. We have also predicted the previously unknown bound pose for CAT. We have illustrated the binding pathways for both ligands detailing which residues interact with the ligands at each step. Our results suggest that CAT can not form critical polar interactions with a tyrosine residue present in the island domain of BRI1, resulting in the lower binding affinity compared to BLD. The attenuated interaction with BRI1 for CAT is due to the absence of an oxygen atom in the B-ring, which is suggested by our long timescale kinetic MC simulations. In addition, we uncover a conserved non-productive binding state for both BLD and CAT, which can potentially decrease the ligand affinity by affecting their on-binding rates. Furthermore, the non-productive pose is more stable for CAT, which may contribute to the difference in bioactivity between BLD and CAT. Finally, we have validated previous studies that illustrates a conformational restructuring of the island domain post ligand binding. Overall, our findings improve the understanding of BRs perception, which allows for future chemical and genetic control of BRI1 receptor activity and the associated biological responses.

## Theoretical Methods

### MD simulations of the binding of BLD and CAT to BRI1

In essence, MD simulations illustrate the motions of the atoms in a simulation system with respect to time.^37^ If the spatial positions of all the atoms at a particular point of time and the net force on each atom acted by the other atoms of the system are known, Newton’s second law is applied to predict the future positions of the atoms.^38^ The MD simulations for both our ligand systems were conducted using the crystal structure of BLD-bound BRI1 as the initial structure (PDB ID: 3RGZ^9^). To save computational time, the receptor protein was truncated from residue 323 to residue 741 using VMD.^39^ It was then capped at the N-terminal with NME and at the C-terminal with ACE. PDBFixer was used to add the missing atoms and hydrogens.^40^ The protonation states of the histidine residues were inspected using H++ server at pH of 7.^41^ The force field used for the protein was AMBER ff14SB^42^ and that for the ligands was the general AMBER force field.^43^ The disulfide bonds formed by the cystine residues were added where necessary using tleap. To neutralize the system and imitate the conditions of the living cells, 150 mM of sodium chloride was added to the system. The ligands were placed at a random position far away from the binding pocket of the BRI1 protein. Both the systems were minimized for 50,000 steps and then equilibrated at 300 K for 10 ns. To capture the long timescale process of ligand binding efficiently in a relatively short amount of time, the strategy of adaptive sampling was employed. ^24–27^ This required orchestrating multiple rounds of simulations, where each round used the configurations of the protein-ligand system generated after the previous rounds as the starting structures for simulations. The configurations were selected according to the configuration density from previous sampling or the distance between the centers of mass of BRI1 and either ligand. All the simulations were run in AMBER14 and AMBER18. ^44^ The simulations were conducted using either a local cluster or the Blue Waters supercomputer. ^45^ The total simulation time was 58 *µ*s for BLD binding and 44 *µ*s for CAT binding (Tables S1 and S2).

### MSM construction and hyperparameter selection

MSMs are powerful tools to analyze large sets of short parallel trajectories, allowing for a rigorous statistical analysis of rare biological events.^28–30^ The MSM approach uses all trajectories to generate a model comprised of a certain number of kinetically relevant states and the transition probabilities between these states.^28–30,46^ To generate an MSM, a selection of features relevant to the system is made. For our ligand-protein systems, these were distance metrics which could accurately describe the movement of the ligands during the binding processes. A set of distances between the atom pairs belonging to BRI1 and both ligands were chosen as the features (Tables S3 and S4). The first subset of features were pairwise distances between the C_*α*_ atoms of the residues within 5 Å of the bound ligands and three atoms in the ligands. The rest of distance features were specific interactions between BRI1 and the bound ligands. Featurization of the simulation datasets was performed using MDTraj 1.7.0. ^47^

Dimensionality reduction was performed for clustering the conformation ensemble from MD simulations. This was implemented using time-lagged independent component analysis (tICA), an approach to analyze the slower dynamics of protein systems.^48^ tICA was performed on the normalized featurization dataset to generate the slowest relaxing degrees of freedom from the linear combinations of the features. Next, the data for each ligand system, representing all the conformations generated from MD simulations, was clustered into a number of states based on several tICs using the *k*-means clustering method. A number of MSMs were then built using the clustering result using different lag times. The lag times, at which the slowest implied timescales captured by these MSMs converge, were chosen for building MSMs (Figure S1). The numbers of clusters and tICs for both systems were further optimized through a cross-validation method which used the Generalized Matrix Rayleigh Quotient (GMRQ) score to estimate the slow dynamic modes of the protein systems.^49^ The numbers of tICs and clusters which corresponded to the highest GMRQ scores for each system, were chosen as the optimum ones to build the MSMs (Figure S2). tICA, clustering, and MSM construction were implemented using MSMBuilder 3.4 python package. ^50^ The iteration processes to estimate GMRQ scores for MSMs were performed using Osprey, which is a tool for hyperparameter optimization of machine learning algorithms.^51^ The graphs were plotted using the Matplotlib 3.1.1 python package.^52^ The final MSM parameters for BLD are 200 clusters, 6 tICs and a lag time of 20 ns, and those for CAT are 80 clusters, 3 tICs and a lag time of 20 ns.

### Estimation of standard binding free energy from potential of mean force

The standard binding free energies (Δ*G*^*°*^) for both BLD and CAT to BRI1 were estimated from the three-dimensional potential of mean force *W* (*r*) (3D PMF).^53,54^ The 3D PMFs were generated by projecting the relative positions of both ligands to BRI1 onto the Euclidean coordinates, weighted by their equilibrium probabilities given by the MSMs (Figures S3 and S4). Δ*G*^*°*^ was then calculated using the following expression:

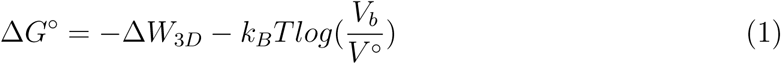

where Δ*W*_3*D*_ is the depth of the 3D PMF, *k*_*B*_ is the Boltzmann constant, T is the temperature, *V*_*b*_ is the bound volume calculated as the integral of the PMF, and *V* ^*°*^ is the standard-state volume, which is 1661 Å^3^. *V*_*b*_ is given by:

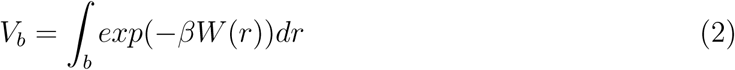

where the bound region is defined by a cutoff distance from the minima of the 3D PMF and a cutoff *W* (*r*). A sensitivity analysis was performed to choose the appropriate cutoff distance and *W* (*r*) to define the bound volume.^54^

### Kinetic MC simulations of ligand binding dynamics

The binding dynamics of the ligands over long timescales can be estimated by kinetic Monte Carlo (MC) simulations. This is essentially a probabilistic approach which relies on the transition probability matrix generated from the MSM models in order to generate long trajectories. Analysis of these trajectories can give kinetically relevant information about important rare biological events captured by the MSMs, such as ligand binding and unbinding.^55^ For both our systems, the starting states were chosen such that the ligands were more than 40 Å away from the receptor. The MSM trajectories were generated using the MSMBuilder 3.4 python package.^50^

### Transition path theory analysis of ligand binding processes

Transition path theory (TPT) serves as a tool to calculate the probabilities and fluxes for the pathways between source and sink states in MSMs. ^56,57^ TPT can be applied in the context of our protein-ligand problems to generate the most probable pathways via which the ligands can reach their final bound states starting from initially unbound states. The unbound states were chosen such that the ligands were more than 30 Å away from the receptor. The bound states were chosen such that the RMSDs of the ligands were less than 2 Å as compared to their respective bound poses and also their distances from S647, an important residue interacting with both ligands in their bound poses, were less than 4 Å. The transition pathways were estimated using the MSMBuilder 3.4 python package. ^50^

### Root mean square fluctuation analysis of the island domain

The flexibility of protein backbone can be captured by calculating the root mean square fluctuation (RMSF) of the C_*α*_ atoms of the residues in BRI1. RMSF is a measure of atomic fluctuations and can be an effective tool to compare differences in protein conformation before and after ligand binding. For both our systems, the RMSF values were generated after superimposing each frame of the trajectories against the conformation of BRI1 bound with BLD. The RMSF calculations were carried out using the Pytraj 2.0.1 python package. ^58^

## Results and Discussion

### Thermodynamic analysis of ligand binding processes reveals a higher affinity for BLD than CAT and key binding intermediate states

Using extensive MD simulations, we captured the binding of both BLD and CAT to the BRI1 receptor. To gain thermodynamic insights into the ligand binding processes, we estimated the standard binding free energy (Δ*G*^*°*^) for both ligands and generated the free energy landscapes that provided the energetic description of the processes. The three-dimensional PMFs along the Euclidean coordinates were produced to estimate the depth of PMF Δ*W*_3*D*_ for BLD and CAT, which are 15.01 *±* 0.12 kcal/mol and 11.02 *±* 0.1 kcal/mol, respectively (Figures S3A-C and S4A-C). We varied the cutoff distance and the cutoff *W* (*r*) to define *V*_*b*_, and then characterized the changes in *V*_*b*_ and Δ*G*^*°*^ (Figures S3D,E and S4D,E). After the sensitivity analysis, we defined the bound conformation as those states with the distance less than 10 Å from the 3D PMF minima and *W* (*r*) less than 2 kcal/mol. The 3D PMFs yielded a standard free energy of binding Δ*G*^*°*^ = -10.77 *±* 0.11 kcal/mol for BLD, and Δ*G*^*°*^ = -6.9 *±* 0.11 kcal/mol for CAT. The error bars on Δ*G*^*°*^ were determined by averaging over 10 PMFs by the projection of random 50% of simulation data, weighted by their equilibrium probabilities generated from Bayesian MSMs. Our predicted Δ*G*^*°*^ for BLD is in good agreement with the experimental 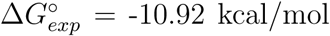.^59^ The lower Δ*G*_*bind*_ for BLD binding suggests that BLD has higher binding affinity as compared to CAT. This is also in accordance with the previous studies reporting that BLD is the most potent BR.^20–23^

To quantitatively characterize the ligand binding processes, we generated the free energy landscapes by projecting the conformations onto two important structural metrics instead of the Euclidean coordinates (Figures 2A,B). For each system, the distance of the ligand from S647 of BRI1 and the RMSD of the ligand from its bound pose were selected as the two structural metrics. The distance from S647 was chosen as this BRI1 residue was observed to interact with the ligands in several intermediate states as well as in their bound states. The RMSD of the ligands from their bound pose was chosen as it accurately reflects the deviation of the ligands from their final bound pose. We also estimated the error bars on the binding free energy landscapes (Figure S5).

**Figure 2:**
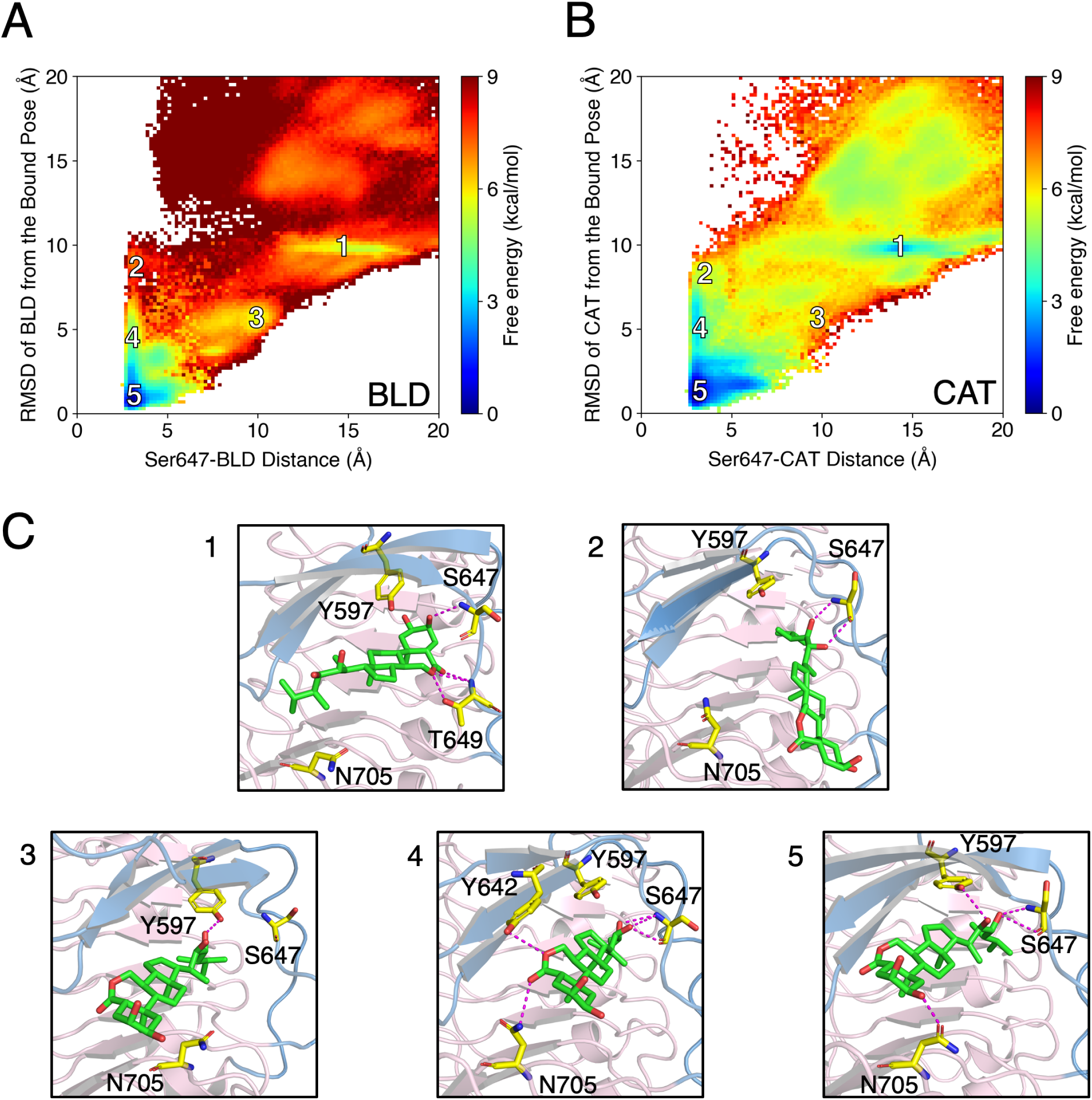
The free energy landscapes for both (A) BLD binding and (B) CAT binding to BRI1 are constructed by projecting all the simulated conformations, weighted by the MSMs, in the context of two metrics including the distance of the ligands from S647 and the RMSD of the ligands from their bound poses. (C) The conformations of the ligand-protein system for BLD corresponding to the state labels in the free energy landscape are shown. The LRR core of BRI1 is shown in light pink and the island domain in sky blue. BLD is shown as green sticks while the important residues of BRI1, involved in direct hydrogen bonding with BLD, are shown as magenta sticks.

A deeper analysis of the free energy landscapes revealed the key conformational states during the binding processes (marked as states 1-5 in Figures 2A,B). The snapshots corresponding to these states for BLD binding are shown in Figure 2C. The states 1-4 are key intermediate states while the state 5 represents the ligand bound configuration of the receptor. In state 1, the rings A-D of BLD are positioned inside the small binding pocket formed by LRRs 21, 22 and the island domain. In this position, the hydroxyl group of C2 of the A-ring interacts with S647 while the oxygens of the B-ring interact with T649. This conformation is flipped with respect to the final bound pose, where the hormone side chain resides in the small binding pocket instead of the fused rings. We thus propose that this might be a non-productive binding pose attained by BLD during binding. This non-productive binding is also captured in the CAT binding process (Figure 2B). These states potentially reduce the affinity of BRs by lowering the on-binding rates. On the other hand in the intermediate state 4, BLD has assumed the orientation in the bound pose but is yet to fix itself in the correct configuration. It interacts with S647 via the vicinal diols of its side chain, with Y642 via the ring oxygen of its B-ring and N705 via the carbonyl oxygen of its B-ring. The positions of the two intermediate states in the free energy landscape indicate that BLD may be able to bind to BRI1 via more than one pathway.

To better show the two different pathways BLD might follow, we characterized states 2 and 3, which appear to precede state 4 and succeed state 1, respectively. State 2 describes a configuration where the BLD side chain appears to approach the binding pocket. The side chain vicinal diols form hydrogen bonding interactions with S647. State 3 corresponds to a configuration where the BLD side chain is oriented to enter the binding pocket but has not done so yet. In this state, BLD forms a hydrogen bonding interaction with the Y597 via the hydroxyl group attached to C23 in its side chain. It is observed from the free energy landscape that there is a considerable energy barrier (*∼*4 kcal/mol) between state 1 and state 3. To reach from state 1 to 3, the linked rings in BLD have to move out of the binding pocket and reverse its orientation to allow the side chain to enter the binding pocket instead. The enhanced stability of the non-productive binding pose described by state 1 may be the reason for such an energy barrier associated with the flipping process. In contrast, the transition from state 2 to state 4 and finally to the bound state 5 faces no significant energy barrier, with the landscape showing gradual decrease in free energy.

In addition, there is a distinct similarity between the BLD and CAT landscapes in terms of the landscape shapes and the positions of the energy minima. Consequently, the intermediate states for the binding of each ligand are also observed to be similar to the degree allowed by their structural difference. An important difference, however, in the case of CAT is the lower free energy associated with the non-productive binding state 1 as compared to BLD (*∼*3 kcal/mol difference). As a result, it is possible that CAT may spend more time in this pose at equilibrium, thus hindering the flipping process to assume the bound pose. This could be another reason for the lower biological potency of CAT. Overall, our thermodynamic results have revealed several key states during ligand binding for both systems and have also provided energetic insights into why BLD is more bioactive than CAT.

### Simulations unveil the previously unknown bound pose for CAT

MD simulations and consequent MSM analysis have captured the bound poses for both BLD and CAT. The pose captured for BLD is identical to that of the crystal structure (Figure 3A) (PDB ID: 3RGZ^9^), with the RMSD of the simulated bound pose less than 1 Å from the crystal structure. We have accurately reproduced several of the BRI1-BLD interactions as well (Figure 3B).^9,12^ The BRI1 residues involved in BLD perception are S647, Y597, Y599 and H645 of the island domain and N705 of the LRR core. Direct hydrogen bonding interactions of BLD with S647 and Y597 via the vicinal diols of its side chain is in agreement with earlier reports. The exact water mediated hydrogen bond interactions have also been predicted correctly.^9,12^ These include three water molecules which mediate interactions between Y599 and oxygens of the B-ring lactone group, S647 and C22 hydroxyl group and finally among H645, Y597 and the C22 and C23 hydroxyl groups (Figures 3B,D). In addition, N705 forms hydrogen bonding interaction with the hydroxyl group of C3 in the A-ring (Figures 3B,D). This demonstrates the ability of MD simulations to accurately illustrate the configuration and interactions of BLD in its bound pose.

**Figure 3:**
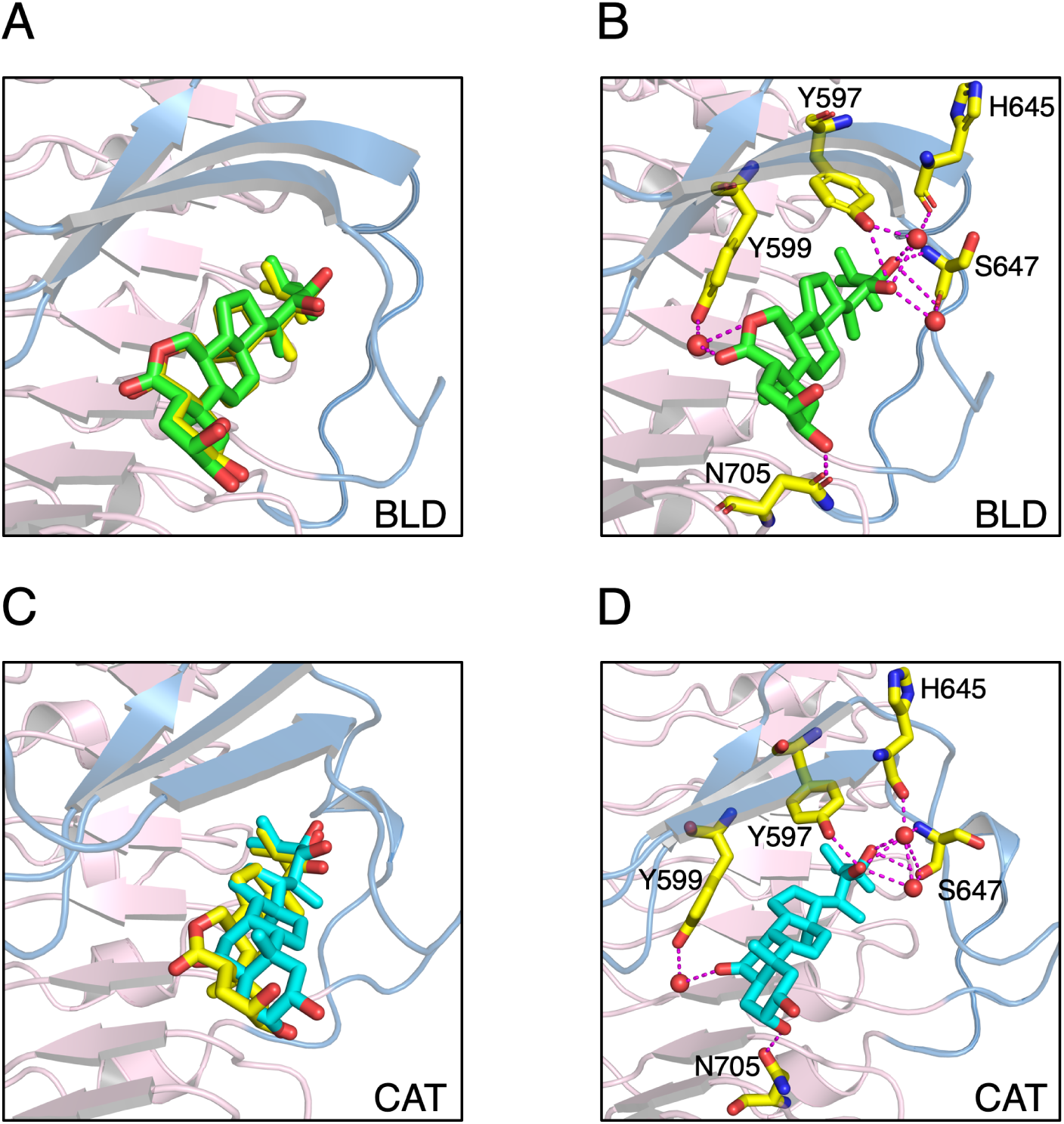
The simulated bound forms of (A) BLD and (C) CAT are superimposed against the BLD-bound BRI1 crystal structure. The important hydrogen bonded interactions, both direct and water-mediated, with the residues of the island domain and LRR core of BRI1 obtained from simulations, are shown for (B) BLD and (D) CAT. BLD corresponding to the crystal structure is shown as yellow sticks, while the BLD and CAT molecules obtained from simulations are shown as green and cyan sticks, respectively.

A crystal structure of CAT-bound BRI1 has yet to be solved. Thus, our simulated bound pose is the first such structure showing the spatial configuration and also the interactions of CAT with BRI1 in the bound state. We report that CAT assumes a similar but a more distorted configuration compared to BLD (Figure 3C). However, it can still interact with the same residues which interact with BLD (Figure 3D). The slightly distorted configuration assumed by CAT is likely due to the absence of the extra oxygen in its B-ring.^19^ We report direct hydrogen bonding interactions of CAT through its side chain hydroxyl groups of C22, C23 and its B-ring C6 attached carbonyl oxygen with Y597, S647 and N705 residues, respectively. We further report water mediated hydrogen bond interactions involving 3 water molecules. They mediate interactions between the hydroxyl group of C22 and S647, carbonyl group of C6 and Y599 and finally among the hydroxyl groups of C22, C23, and, H645 and S647. Therefore, we have captured the bound pose of CAT in atomic level detail, which provides structural insights into the lower affinity for CAT.

### BLD shows greater stability in the bound pose than CAT

The binding dynamics of the two ligands over long timescales was illustrated using kinetic Monte Carlo simulations on the MSM transition probability matrix. We generated 50 *µ*s long trajectories showing binding and unbinding of the ligands (Figure 4). For both the ligand systems, we selected two specific residues of BRI1, whose distances from the ligand, observed over time, could represent the binding dynamics accurately. The distance between S647 and the hydroxyl group in the side chain of each ligand was chosen to represent the binding of the ligands and also the configuration changes of the side chain upon binding (Figure 4A). The distance between Y599 and the B-ring carbonyl oxygen of the ligand was chosen to represent the spatial fluctuations of the fused rings at the other end of the ligand (Figure 4B). For both systems, the average distance of the ligand side chain from the serine residue is *∼*3 Å when bound, indicating that the ligands position themselves at similar depths inside the binding pocket of BRI1. On the other hand, the average distance of the tyrosine residue from the bound BLD is *∼*5 Å as compared to *∼*3 Å from the bound CAT. This difference results from the extra oxygen in the B-ring of BLD which pushes the interacting carbonyl oxygen further away from the tyrosine. Our results show a higher degree of ligand fluctuations for CAT compared to BLD, especially highlighted by its interaction with Y599 (Figure 4B). We propose that this difference arises from the absence of the hydrogen bonding interaction between Y599 and the non-existent B-ring oxygen in CAT (Figures 3B,D). This results in the lower binding stability for CAT, and may thus be a major reason why CAT was found to be less bioactive than BLD in previous research.^20–23^ Overall, our study reflects that BLD is more stable than CAT on BRI1 binding, primarily due to the presence of an extra oxygen in the BLD.

**Figure 4:**
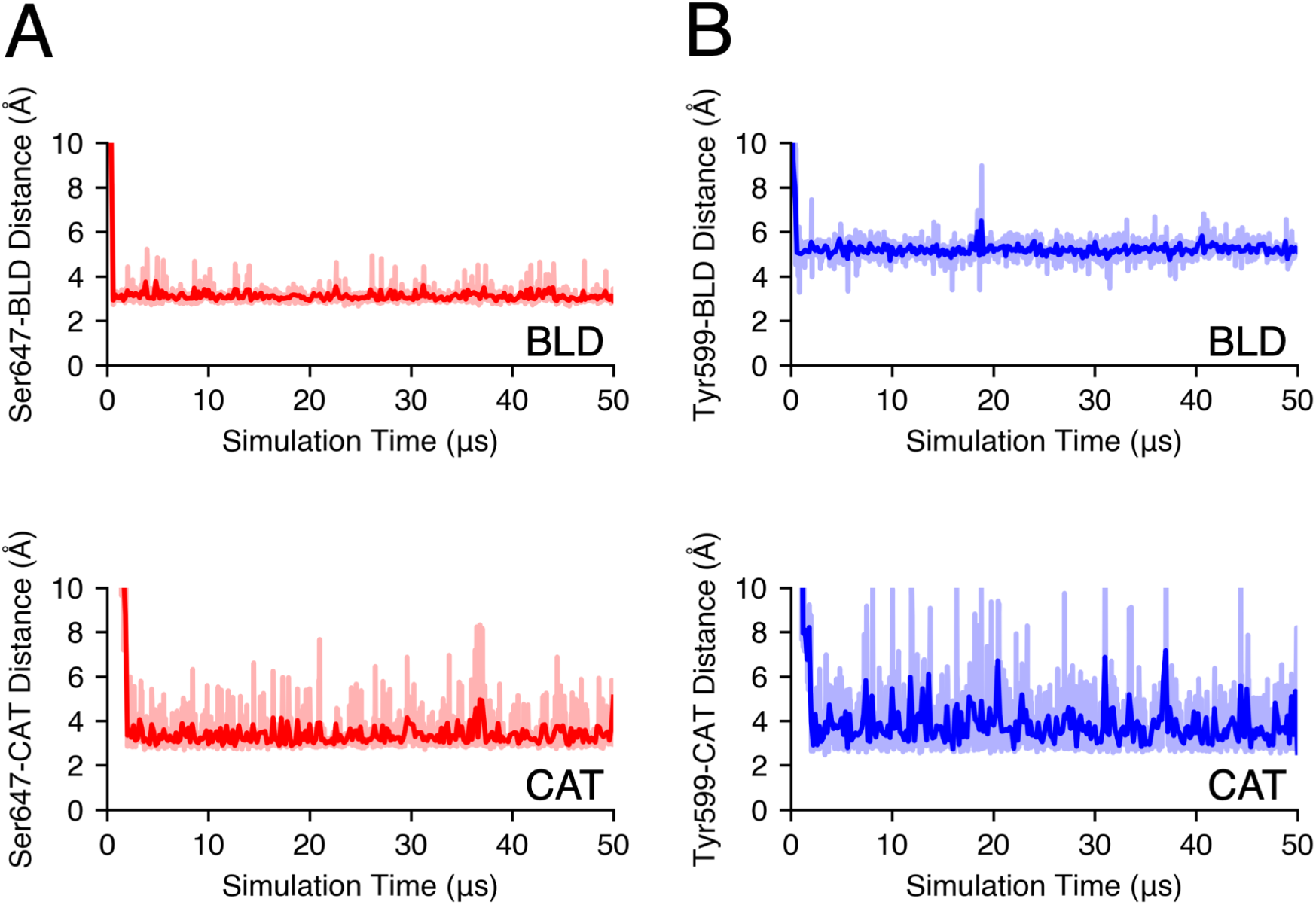
Kinetic Monte Carlo simulations were performed to capture the binding dynamics of both ligands over long timescales. The total simulation time is 50 *µ*s. (A) The distance between S647 and the side chain of either ligand is chosen to represent the binding of the ligands to the receptor and the configuration changes of the side chain upon binding. (B) The distance between Y599 and the B-ring carbonyl oxygen of either ligand is chosen to represent the spatial fluctuations of the fused A-D rings at the other end of the ligand. CAT shows a higher degree of spatial fluctuations than BLD, indicating lower stability in the bound pose.

### Tansition path theory reveals similar binding mechanisms for BLD and CAT

To analyze the binding pathways for BLD and CAT in high resolution, we applied the transition path theory (TPT) to the MSMs and obtained the top favorable pathways for ligand binding along with their reactive fluxes.^56,57^ For both systems, we generated binding pathways starting from the MSM states where the ligands were more than 30 Å away from BRI1. We report the top two major pathways for BLD and CAT (Figures 5 and 6). For both cases, the top two pathways have similar fluxes. In the first pathway for BLD binding (path I in Figure 5), BLD is observed to approach the BRI1 binding pocket with its fused A-D rings facing the binding groove. It then proceeds further into the pocket to interact with S647 of the BRI1 island domain via its hydroxyl group in the A-ring C2. Next, the hormone rings appear to move away from the binding pocket. This may represent a part of the flipping process where BLD reverses orientation to allow its side chain to enter into the pocket and ultimately assume the bound pose. At this stage, it still maintains the previous interaction with S647, but develops new interactions with T649 via the two oxygens in its B-ring. Finally, BLD assumes its bound pose. The second binding pathway followed by BLD (path II in Figure 5) stands in contrast to the previous one in the sense that instead of the fused rings of BLD, now the side chain of the ligand leads it towards the binding pocket. In this first state, the hydroxyl groups of C22 and C23 of the side chain interact with S647, and BLD has not entered the BRI1 binding groove yet. Thereafter, BLD enters the binding pocket but is still not in its bound configuration. It continues the previous interaction with S647, meanwhile forming new interactions with Y642 and N705 through the ring oxygen and the carbonyl oxygen of its B-ring, respectively. Finally, BLD binds to the BRI1 receptor. Thus, we have captured two distinct binding pathways of BLD with BRI1, confirming the results from the free energy landscapes.

**Figure 5:**
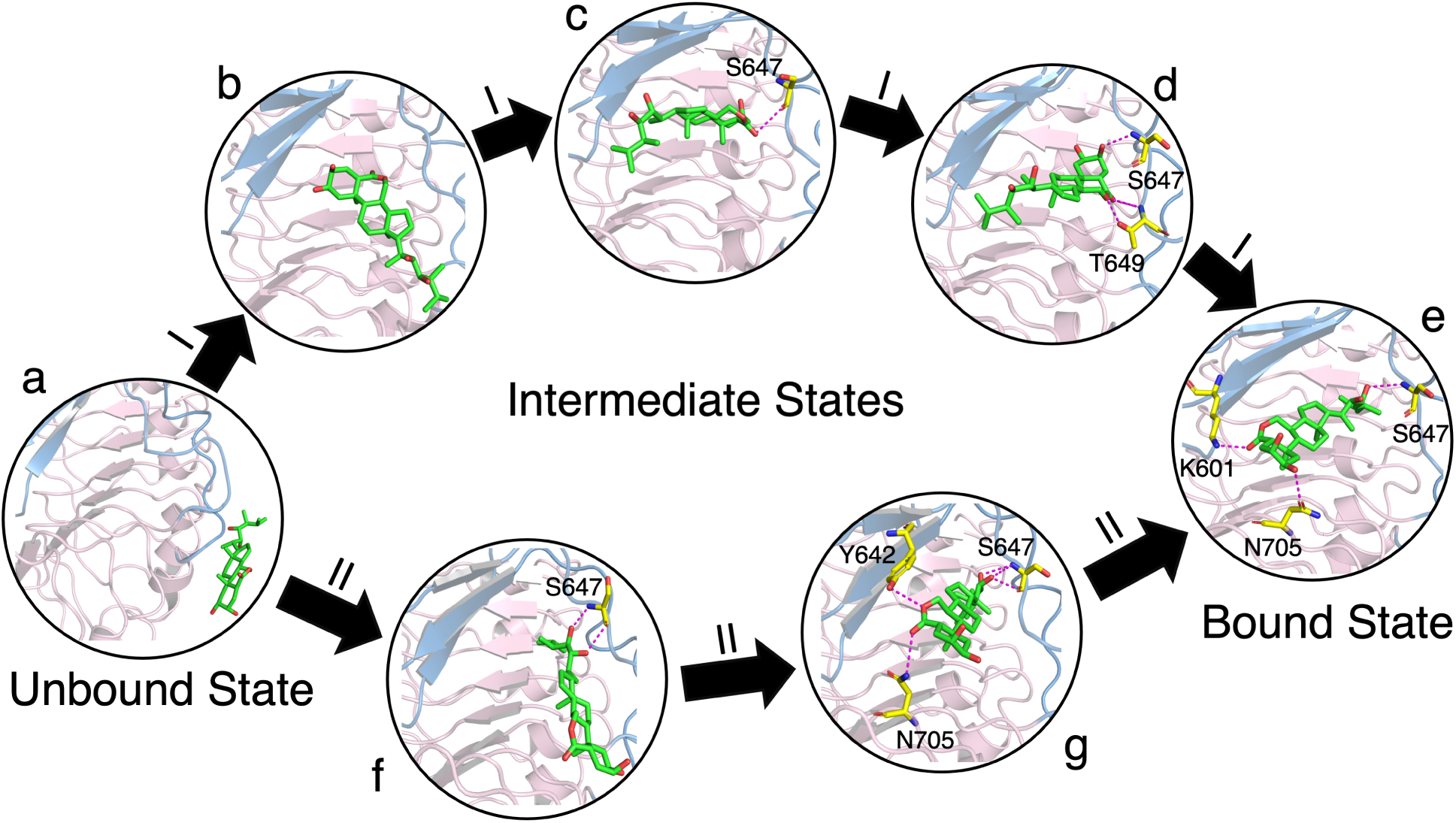
The top two pathways of BLD binding to BRI1 obtained using transition path theory have similar fluxes. The unbound state is represented by (a) and the bound state by (e). Path I comprises of the states (a), (b), (c), (d) and (e), and outlines a binding pathway where the A-D rings of BLD enters the BRI1 binding pocket first and then flips orientation to allow the side chain to enter the binding pocket instead, before finally assuming the bound pose. Path II comprises of the states (a), (f), (g) and (e), and represent a binding pathway where the BLD side chain leads the ligand into the binding pocket and gradually positions itself in the correct bound pose.

**Figure 6:**
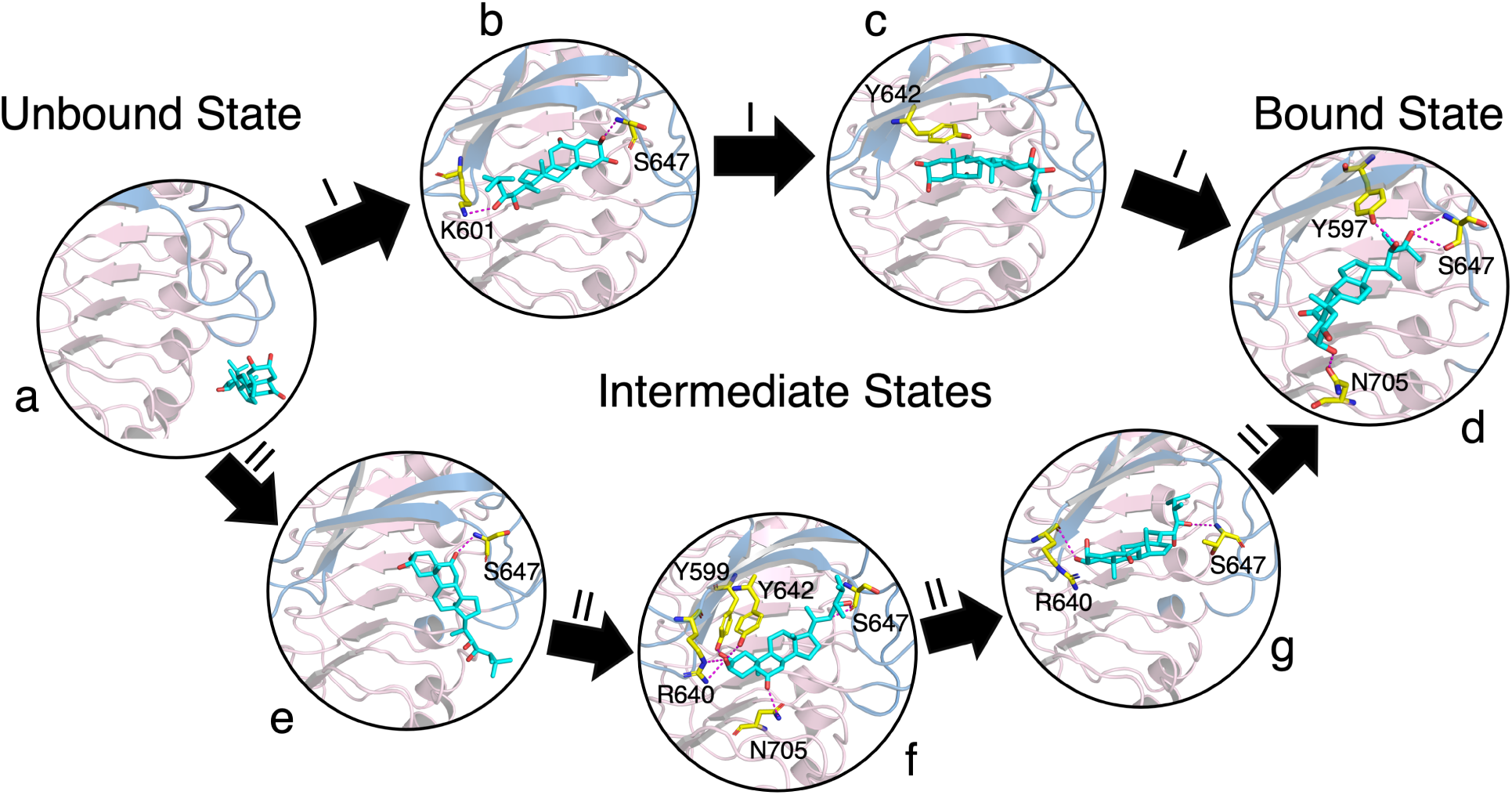
The top two pathway of CAT binding to BRI1 obtained using using transition path theory have similar fluxes. The unbound state is represented by (a) and the bound state by (d). Path I comprises of the states (a), (b), (c) and (d), and outlines a binding pathway where the A-D rings of CAT enters the BRI1 binding pocket first and then flips orientation to allow the side chain to enter the binding pocket instead, before finally assuming the bound pose. Path II comprises of the states (a), (e), (f), (g) and (d), and represent a binding pathway where the CAT side chain appears to lead the entire ligand into the binding pocket before gradually positioning itself in the correct bound pose.

TPT analysis further generated two favorable pathways for CAT binding. The pathways captured for CAT suggests that it approaches BRI1 in a similar fashion as BLD. The first pathway for CAT binding (path I in Figure 6) is similar to path I for BLD binding in the way that the fused A-D rings of CAT enter the BRI1 binding pocket first instead of the side chain. In the first step of this pathway, hydrogen bonding interactions are formed by the hydroxyl groups of C3 in the A-ring and C23 in the side chain with S647 and K601, respectively. Following this, the fused rings move out of the BRI1 binding pocket and form a ring-ring hydrophobic interaction with Y642. The transition to this state resembles a similar flipping of the ring group as observed in the case of BLD. Finally, CAT enters the binding pocket and assumes its bound pose. The second pathway for CAT binding (path II in Figure 6) starts with the B-ring carbonyl oxygen forming a hydrogen bond with S647. CAT has not yet entered the binding pocket at this stage. Following this, the side chain of CAT leads it towards the binding groove. In this state, the B-ring carbonyl oxygen and the side chain hydroxyl groups form hydrogen bonds with N705 and S647, respectively. Further, the hydroxyl group of C2 in the CAT A-ring forms a hydrogen bond with Y599, while the hydroxyl group of C3 beside it forms hydrogen bonds with R640 and Y642. Thereafter, CAT assumes a configuration where it is deeper inside the binding pocket but not yet bound. In this state, the hydroxyl groups of the A-ring C2 and the side chain C23 form hydrogen bonding interactions with R640 and S647, respectively. Finally, CAT positions itself perfectly to assume the bound pose. This second pathway is similar to path II of BLD binding as it appears that the side chain leads the ligand into the binding pocket here as well. Overall, our results have characterized two distinct binding pathways of both BLD and CAT and have reported similarities in their binding mechanisms.

### BRI1 island domain undergoes a major conformational restructuring upon ligand binding

To quantitatively analyze the effect of ligand binding on the BRI1 island domain conformation, we calculated the root mean square fluctuations (RMSF) of the C_*α*_ atoms of all BRI1 residues. The island domain was reported to undergo a major conformational change upon the binding of brassinolide by previous studies.^9,12^ RMSF values capture the fluctuations of the residues of a protein domain, with lower RMSF values indicating higher conformational stability. For our ligand systems, the ligand unbound and bound states were used to calculate the fluctuations of the BRI1 residues before and after ligand binding, respectively. We further generated the free energy landscapes for both BLD and CAT projected onto the RMSDs of the island domain and the ligand compared with their corresponding bound poses. The RMSF values of BRI1 residues and the free energy landscape for BLD-BRI1 system are shown in Figures S6A,B and those for BRI1-CAT system are shown in Figures S6A,E respectively. A visual illustration of the protein fluctuations due to ligand binding is shown in Figures S6C,D and S6F,G for BLD and CAT, respectively. We report a significant reduction in RMSF values of the residues of the BRI1 island domain after BLD binding. This result is further supported by the free energy landscape which indicates stable bound states for deviations of the island domain less than 2 Å from the bound crystal structure. Thus, we validate previous studies which report that BLD binding induces the island domain of BRI1 to become more ordered and fixed with respect to the LRR core.^9,12^ We further report a similar stabilization of the island domain upon CAT binding. However, the reduction in RMSF values for the island domain residues (notably residues 640-650) when CAT binds is relatively less compared to BLD. These results suggest that CAT induces a slightly lower degree of stabilization of the island domain, which may in turn increase the ligand off-binding rate. This may, thus, be another reason as to why CAT is less biologically active than BLD. Furthermore, the island domain is involved in BRI1 association with BAK1,^9,12^ and the lower degree of stabilization of island domain may also affect the BRI1-BAK1 association. Overall, our results suggest that both BLD and CAT induces an ordering of the island domain, but that this is a more prominent feature in BLD binding.

## Conclusions

The perception of BRs by the extracellular domain of the plant receptor BRI1 is a key step in brassionsteroid signaling. However, knowledge about the perception mechanism is limited due to the availability of only two crystal structures for the BLD-bound BRI1. ^9,12^ Importantly, it is unclear why major bioactivity differences exist between BRs with strikingly similar structures, which is the most highlighted by the activity difference between BLD and CAT.^20–23^ Using long timescale MD simulations and MSM analysis of the massive ligand binding trajectories, we have provided energetic, dynamic, and structural insights into the binding mechanisms of both BLD and CAT. In addition, we have provided several explanations for the difference in bioactivity as shown by BLD and CAT.

Our results have provided a thermodynamic basis for the perception of BRs and the bioactivity difference between BLD and CAT. From three-dimensional potential of mean force, we have determined the standard binding free energy for both BLD and CAT, corresponding to -10.77 *±* 0.11 kcal/mol for BLD and -6.9 *±* 0.11 kcal/mol for CAT respectively. The predicted Δ*G*^*°*^ for BLD closely matches with the experimental 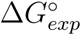 which is -10.92 kcal/mol.^59^ Our free energy calculation results suggest that BLD has relatively higher binding affinity to BRI1 than CAT (*∼*3.9 kcal/mol difference), which is in agreement with the reported bioactivity for BLD and CAT.^20–23^ We have also accurately replicated the bound pose for BLD as captured in the crystal structures. More importantly, we have unveiled the previously unknown bound pose for CAT, which is distorted outward the pocket compared to BLD. The discovery of CAT bound pose allows for structural insights into the origin of its lower binding affinity. We propose that the lower binding affinity arises because CAT lacks an oxygen in its B-ring which forms an important hydrogen bonding interaction with a tyrosine residue of the BRI1 island domain. This is further supported by the kinetic Monte Carlo simulation results that CAT demonstrates significantly higher spatial fluctuations at the fused A-D rings side as compared to BLD. In addition, we propose that the distorted CAT bound pose may attenuate BRI1 association with BAK1 compared to BLD, since the bound BRs directly interact with BAK1 through two hydroxyl groups in the A-ring.^9,12^ Overall, our results suggest that the absence of an oxygen in CAT may be the major reason for the lower bioactivity of CAT as compared to BLD.

Our free energy landscapes and transition path theory analysis on the MSMs have captured the key intermediate states in the ligand binding pathways for both BLD and CAT. We report two distinct pathways with similar fluxes for either ligand. The difference in these two pathways lies in the way the hormones approach the BRI1 binding pocket. We report that either the linked A-D rings of the ligands enters the binding pocket first and later flips orientation to allow the side chain to enter the pocket, or the side chain enters the binding pocket first and then the ligands gradually fix themselves in their final bound poses. We have further captured the BRI1 residues interacting with the two ligands during binding. These residues are a part of either the island domain or the surrounding LRR core, thus confirming the hypothesized importance of these BRI1 regions in BR binding.^9,12^ Thus, we have characterized the binding pathways followed by BLD and CAT in atomistic detail, which provides structural insights into the binding process of BRs.

We report a non-productive binding pose for both BLD and CAT and reveal that this pose is more stable for the latter. It is likely that this intermediate state hinders both ligands from assuming the correct bound pose, thereby lowering the affinity by decreasing the ligand on-binding rates. Since the non-productive binding pose for CAT is *∼*3 kcal/mol more stable as compared to that for BLD, it may further contribute to the lower bioactivity of CAT. Interestingly, similar non-productive binding poses are observed for the binding plant hormone abscisic acid to both PYL5 and PYL10 receptors in *Arabidopsis thaliana*.^31^ Overall, the discovery of non-productive binding poses may provide some guidance on future agrochemical discovery to design ligands with enhanced affinities for targeting these plant receptors.

Our results have also confirmed the hypotheses given by past studies that the island domain undergoes a major structural restructuring upon ligand binding.^9,12^ We report a significant reduction in RMSF values for the island domain residues once the ligands bind to the BRI1 receptor. This is further confirmed by free energy landscapes generated using the RMSDs of the ligand and the island domain from the ligand bound pose. We infer from the landscapes that the stable ligand bound states are the ones which show a higher degree of fixing of the island domain with respect to the BRI1 LRR core. These results are true for both ligand systems, indicating that the binding of BRs stabilizes the island domain in general. However, we observed that CAT binding results in a smaller reduction in RMSF values of the island domain compared to BLD. Since the island domain is directly involved in interaction with both BRs and BAK1,^9,12^ we thus propose that the reduced conformational restructuring of the island domain on CAT binding might be an additional reason for the lower bioactivity of CAT compared to BLD. Overall, our results confirm previous studies that conformational restructuring of the BRI1 island domain is an important event in BR perception.

In conclusion, our study provides critical molecular insights into the binding of the two most potent BR, BLD and CAT, by BRI1. A detailed idea of the binding pathways presented here may allow for a more directed study into BRs perception in plant cells. The results probing the differences in binding of the two ligands as a result of their structural dissimilarities may help future research on synthesizing more active BR analogs. Finally, the leucine-rich repeat receptor-like kinases in plants play a vital role in the perception of various plant peptides and further modulate plant growth, development and stress responses. ^60,61^ Exporting the computational methods used in this study to gain detailed atomistic pictures of the binding of plant peptides and other phytohormones, to their respective receptors, might lead to a better understanding of plant signaling mechanisms.

## Supporting Information Available

Supporting tables and figures. This material is available free of charge via the Internet at http://pubs.acs.org.

## Acknowledgement

The authors acknowledge the support from the Blue Waters sustained-petascale computing project, which is funded by the National Science Foundation (awards OCI-0725070 and ACI-1238993) and the state of Illinois. F.A. acknowledges the support by the Blue Waters Student Internship Program funded by the Shodor Education Foundation. A.D. acknowledges the support by the Khorana Program for Scholars funded by the Department of Biotechnology, Government of India. D.S. acknowledges the support from Foundation for Food and Agriculture Research via the New Innovator Award in Food & Agriculture Research.

## Graphical TOC Entry

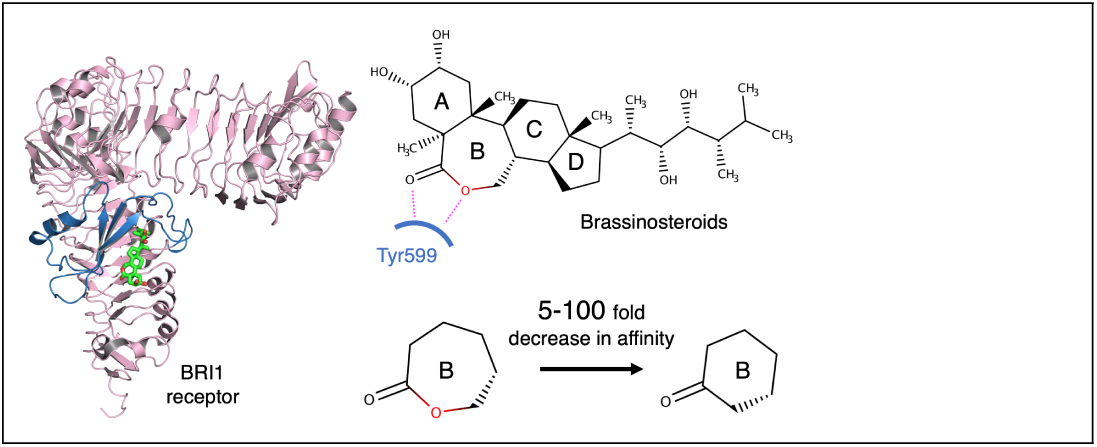

