## Supplemental Information for "Molecular Mechanism of Brassinosteroids Perception by the Plant Growth Receptor BRI1"

### Supporting Information for: Molecular Mechanism of Brassinosteroids Perception by the Plant Growth Receptor BRI1

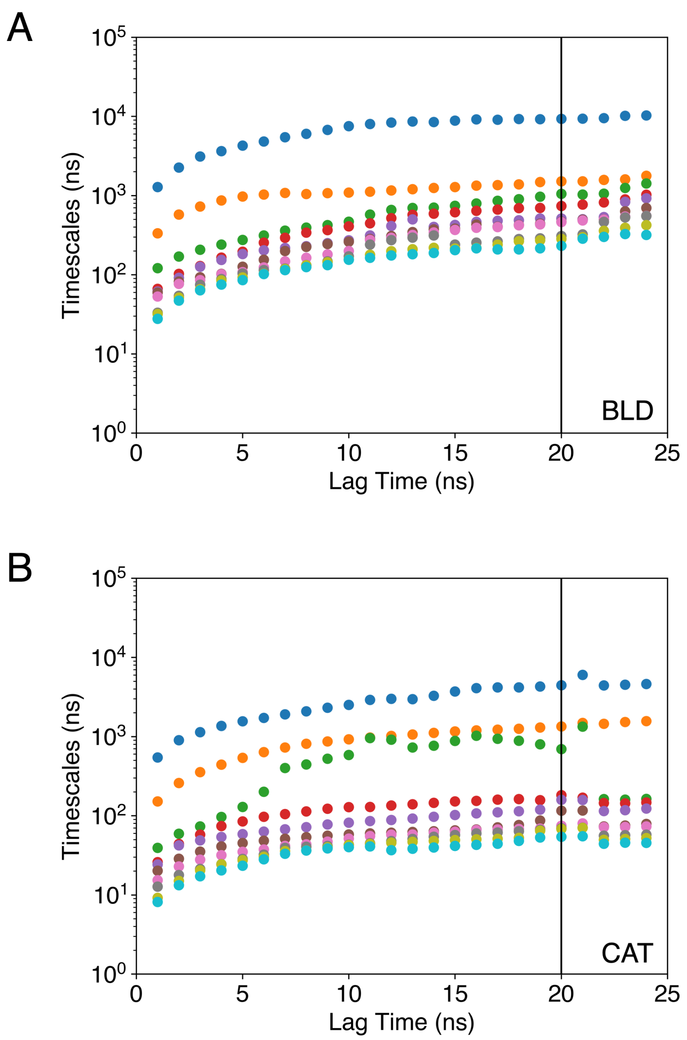


Figure S1: The implied timescales obtained from the transition probability matrix of the (A) 200 state MSM model for BLD and the (B) 80 state MSM model for CAT are plotted against different lag times. Convergence of the implied timescales at a particular lag time implies that the MSM models become Markovian (memoryless) at that lag time. For both systems, convergence is observed at a lag time of 20 ns.

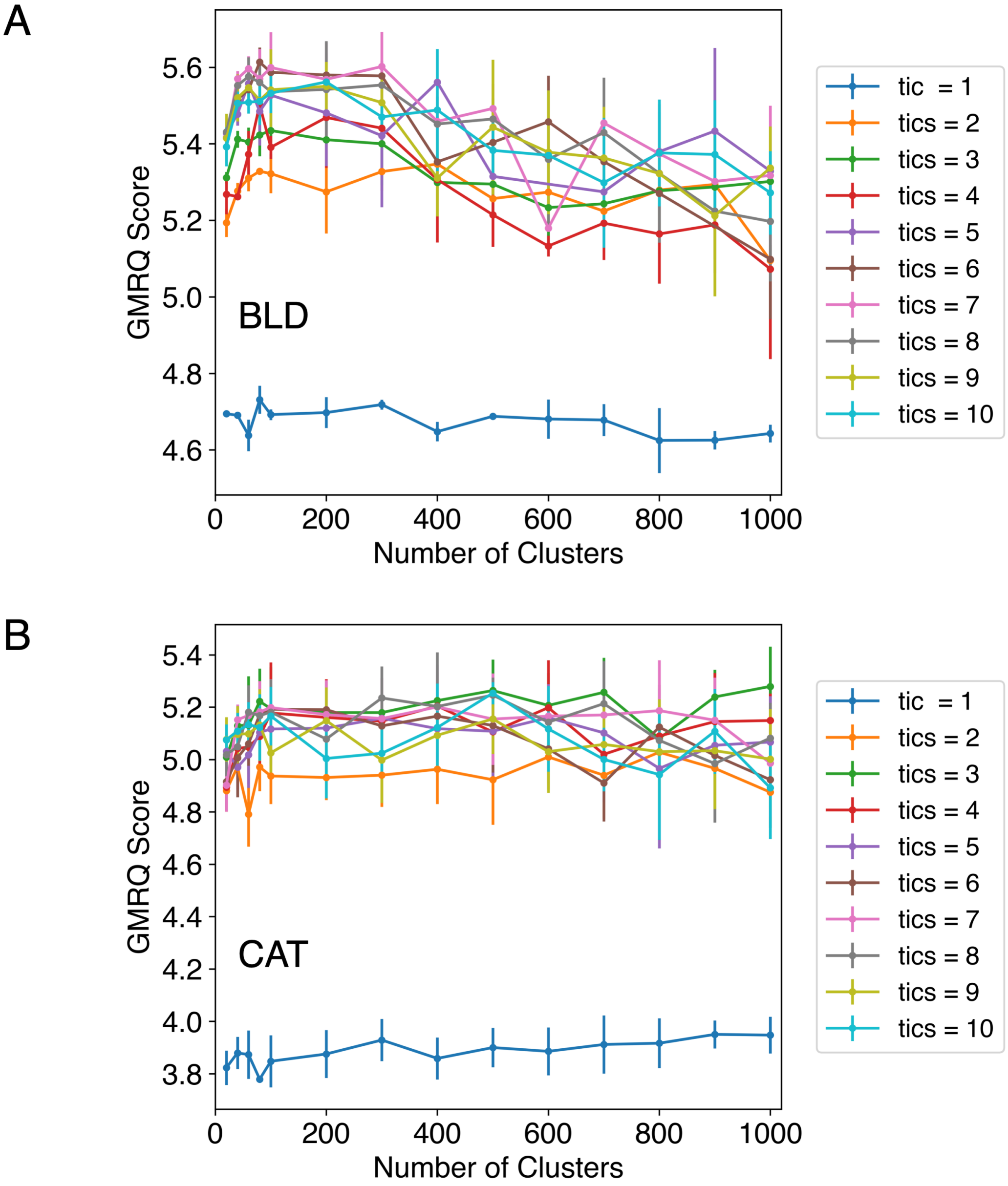


Figure S2: GMRQ scores were generated for a variable number of clusters and tICs for both the ligand systems. The error bars represent the standard deviation from the mean. The highest scores represent the best parameters to build MSMs. The parameters chosen for the (A) BLD-BRI1 system are 200 clusters and 6 tICs and that for the (B) CAT-BRI1 system are 80 clusters and 3 tICs. They were taken as the ideal parameters based on higher scores and good connectivity of the MSM models generated.

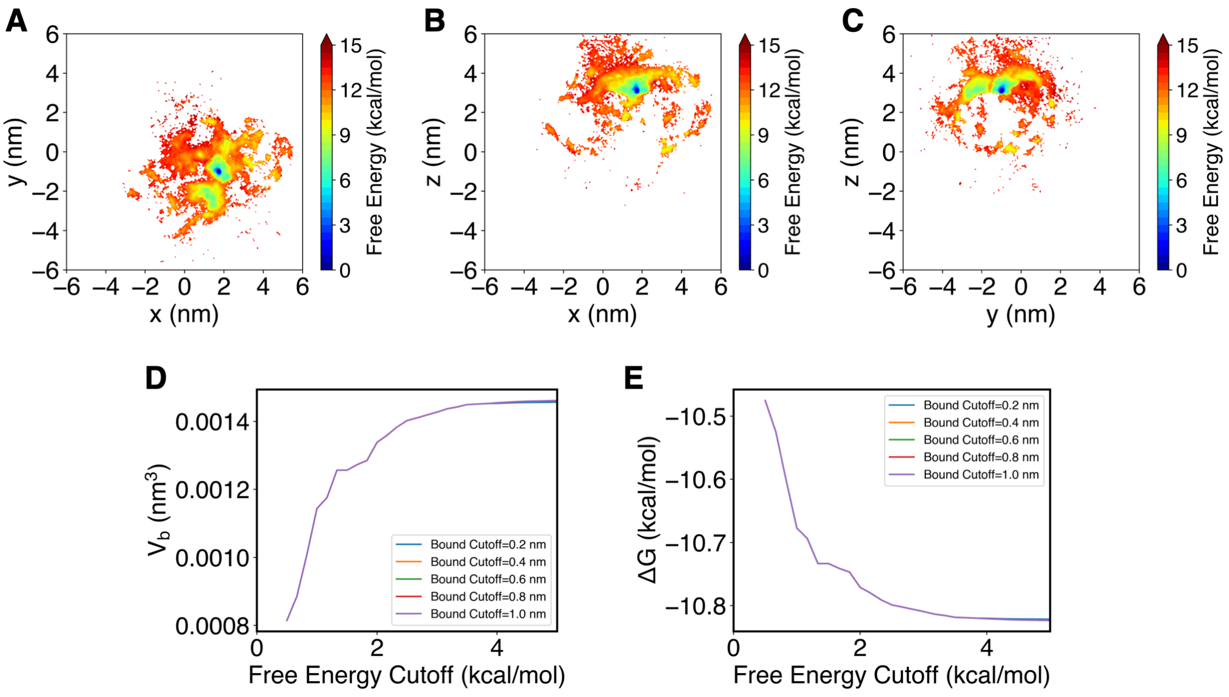


Figure S3: Free energy landscapes of BLD binding to BRI1 projected onto (A) the xy plane, (B) the xz plane and (C) the yz plane. (D) Analysis of the sensitivity of the bound volume ($V_b$) with respect to the cutoff distance and free energy from the minima that are used for the definition of the bound state. (E) Analysis of the sensitivity of the standard binding free energy with respect to the definition of bound volume.

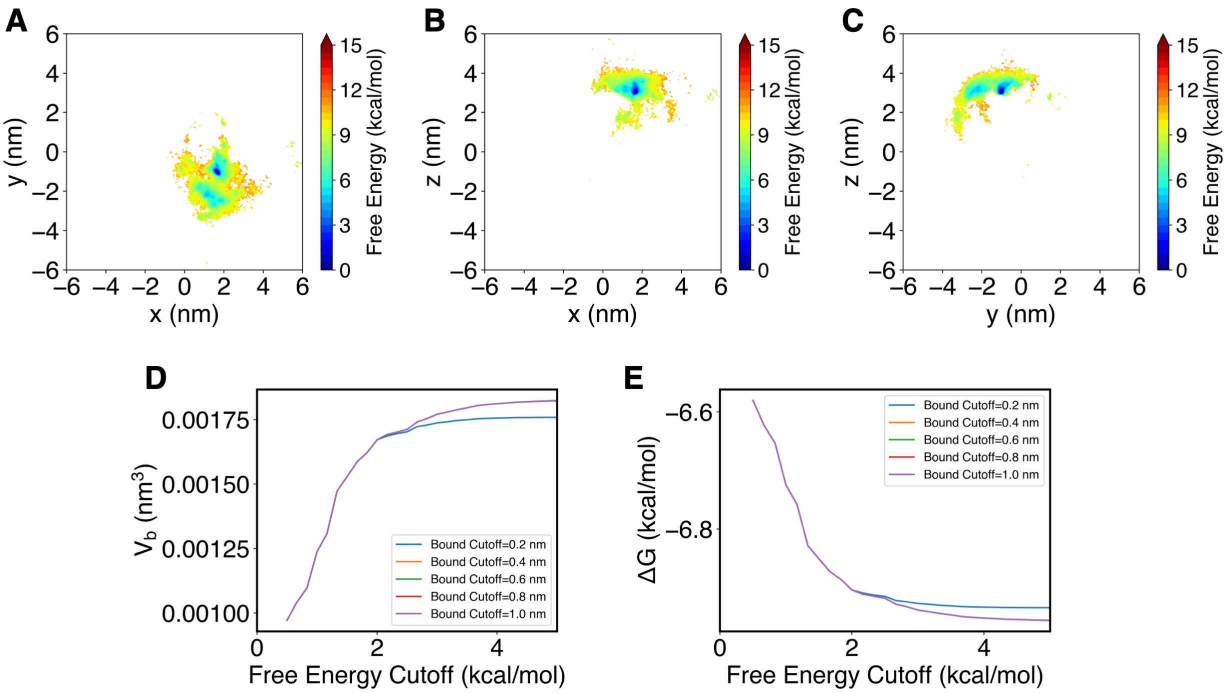


Figure S4: Free energy landscapes of CAT binding to BRI1 projected onto (A) the $xy$ plane, (B) the $xz$ plane and (C) the $yz$ plane. (D) Analysis of the sensitivity of the bound volume ($V_b$) with respect to the cutoff distance and free energy from the minima that are used for the definition of the bound state. (E) Analysis of the sensitivity of the standard binding free energy with respect to the definition of bound volume.

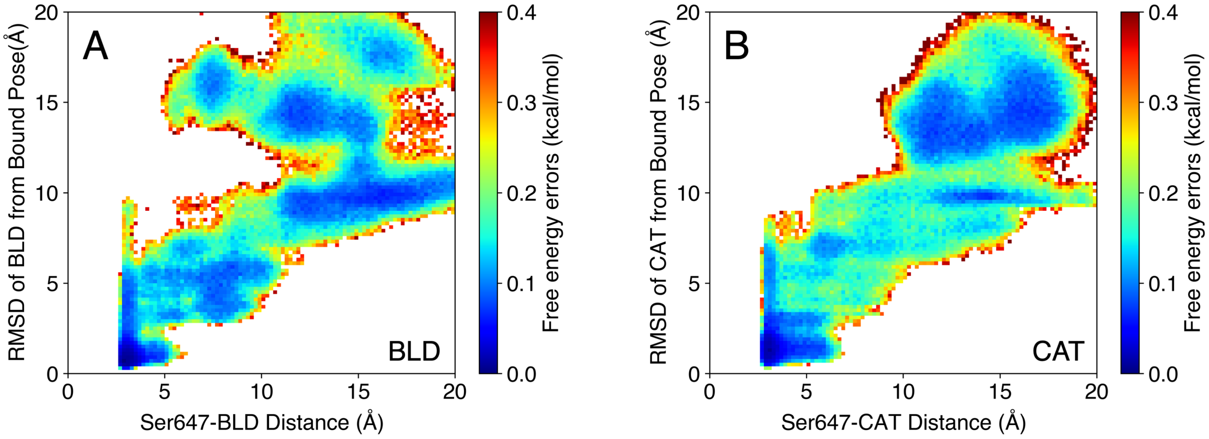


Figure S5: Error bars for the free energy landscape (Figure 2) were obtained by implementing Bayesian Markov State Models over 100 samples and then projecting random 50% simulation data to obtain weighted landscapes in terms of the structural metrics. The standard deviations of the free energies in the landscapes were determined as the error bars.

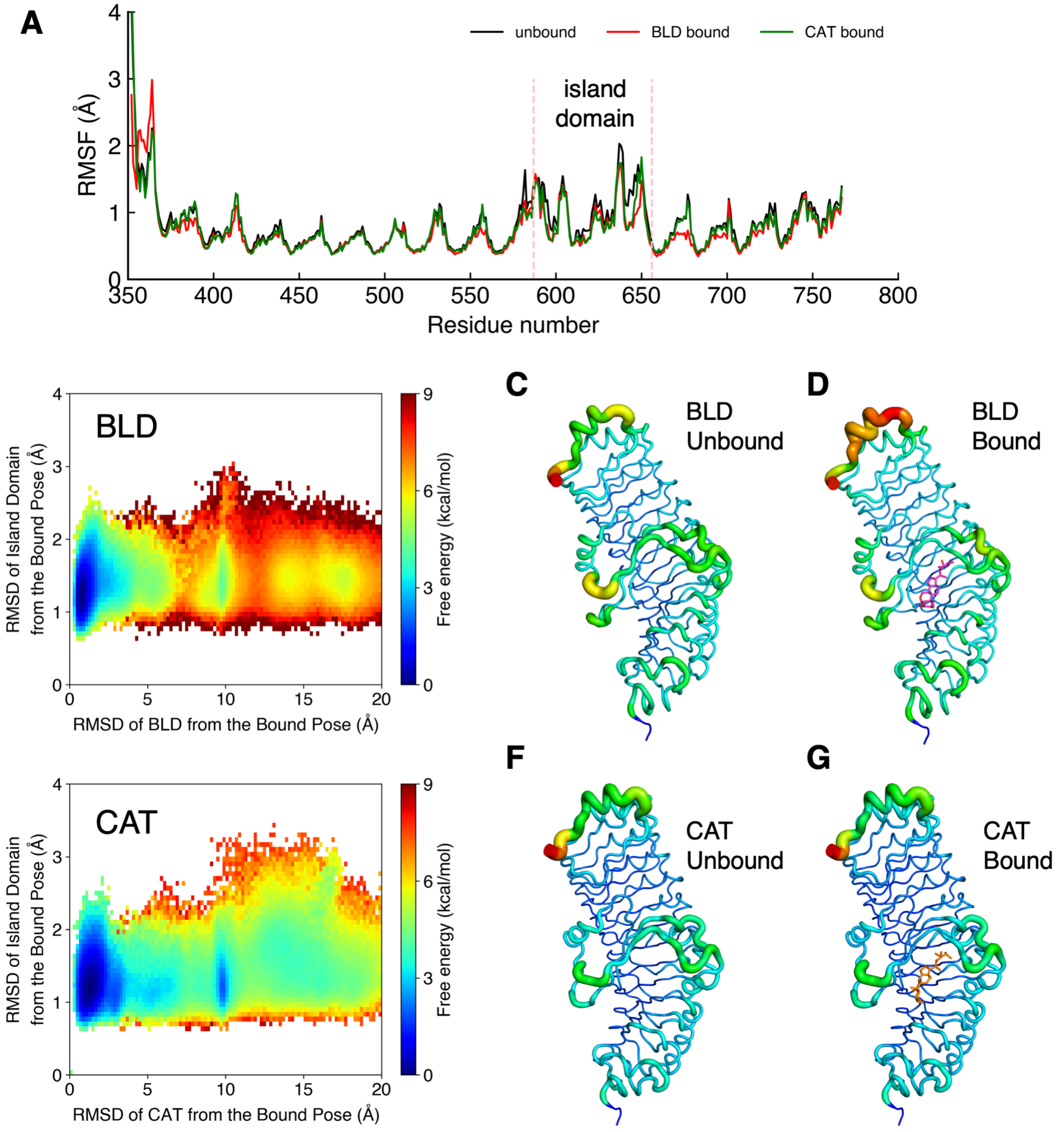


Figure S6: (A) The root mean squared fluctuations (RMSF) was calculated for the $C_\alpha$ atoms of BRI1 residues before and after BLD/CAT binding. The residues making up the island domain show reduction in their RMSF values when BLD/CAT is bound. (B) and (E) The free energy landscapes generated by using the RMSDs of the island domain and BLD/CAT from the bound crystal structure as the two metrics. (C) and (D) Visualization of the RMSF values for the BRI1 residues when BLD is unbound and bound. (F) and (G) Visualization of the RMSF values for the BRI1 residues when CAT is unbound and bound. The tube thickness of the ribbons indicates the fluctuations of the residues, with higher thickness of a protein region corresponding to higher fluctuations of the residues making up that region.

Table S1: Summary of MD simulations of BLD binding to the BRI1 receptor.

| Round | Parallel simulations | Simulation time (ns) | Aggregate ($\mu$s) |
|---|---|---|---|
| 1 | 20 | 58 | 1.16 |
| 2 | 20 | 58 | 1.16 |
| 3 | 20 | 58 | 1.16 |
| 4 | 20 | 58 | 1.16 |
| 5 | 20 | 58 | 1.16 |
| 6 | 20 | 58 | 1.16 |
| 7 | 20 | 58 | 1.16 |
| 8 | 20 | 58 | 1.16 |
| 9 | 20 | 58 | 1.16 |
| 10 | 20 | 58 | 1.16 |
| 11 | 10 | 58 | 0.58 |
| 12 | 10 | 58 | 0.58 |
| 13 | 10 | 58 | 0.58 |
| 14 | 10 | 58 | 0.58 |
| 15 | 10 | 58 | 0.58 |
| 16 | 10 | 58 | 0.58 |
| 17 | 10 | 58 | 0.58 |
| 18 | 50 | 58 | 2.9 |
| 19 | 100 | 58 | 5.8 |
| 20 | 100 | 58 | 5.8 |
| 21 | 60 | 58 | 3.48 |
| 22 | 100 | 58 | 5.8 |
| 23 | 100 | 58 | 5.8 |
| 24 | 220 | 58 | 12.76 |
| Total simulation time: $\sim$58 $\mu$s | | | |

Table S2: Summary of MD simulations of CAT binding to the BRI1 receptor.

| Round | Parallel simulations | Simulation time (ns) | Aggregate ($\mu$s) |
|---|---|---|---|
| 1 | 20 | 60 | 1.2 |
| 2 | 49 | 60 | 2.94 |
| 3 | 50 | 60 | 3.0 |
| 4 | 50 | 60 | 3.0 |
| 5 | 50 | 60 | 3.0 |
| 6 | 20 | 60 | 1.2 |
| 7 | 98 | 60 | 5.88 |
| 8 | 50 | 60 | 3.0 |
| 9 | 100 | 60 | 6.0 |
| 10 | 250 | 60 | 15 |
| Total simulation time: $\sim$44 $\mu$s | | | |

Table S3: Featurization metrics used for analyzing the BRI1-BLD simulation datasets. In total, 80 distances between the atoms in BRI1 and BLD are calculated, including 72 distances that describe the ligand position and 8 distances that describe specific protein-ligand interactions when the ligand is bound.

| BLD position **(72 distances)** | | | |
|---|---|---|---|
| | BLD:O2 | BLD:O23 | BLD:C8 |
| A539:CA<br>I540:CA<br>I563:CA<br>W564:CA<br>I592:CA<br>Y597:CA<br>Y599:CA<br>K601:CA<br>L615:CA<br>Y642:CA<br>G643:CA<br>H645:CA<br>T646:CA<br>S647:CA<br>P648:CA<br>T649:CA<br>M657:CA<br>F658:CA<br>F681:CA<br>I682:CA<br>N705:CA<br>I706:CA<br>M727:CA<br>T729:CA | 1-24 | 25-48 | 49-72 |
| BLD-BRI1 interaction **(8 distances)** | | | |
| Y597:OH | BLD:O22 | 73 | |
| S647:O | BLD:O22 | 74 | |
| Y597:OH | BLD:O23 | 75 | |
| S647:N | BLD:O23 | 76 | |
| S647:O | BLD:O23 | 77 | |
| K601:NZ | BLD:O06 | 78 | |
| N705:OD1 | BLD:O02 | 79 | |
| Y642:OH | BLD:O03 | 80 | |

Table S4: Featurization metrics used for analyzing the BRI1-CAT simulation datasets. In total, 92 distances between the atoms in BRI1 and CAT are calculated, including 75 distances that describe the ligand position and 17 distances that describe protein-ligand interactions when the ligand is bound.

| CAT position **(75 distances)** | | | |
|---|---|---|---|
| | CAT:O2 | CAT:O23 | CAT:C8 |
| L541:CA<br>W564:CA<br>L565:CA<br>A593:CA<br>G594:CA<br>V598:CA<br>I600:CA<br>N602:CA<br>L616:CA<br>V641:CA<br>G643:CA<br>G644:CA<br>T646:CA<br>S647:CA<br>P648:CA<br>T649:CA<br>F650:CA<br>F658:CA<br>L659:CA<br>I682:CA<br>L683:CA<br>I706:CA<br>L707:CA<br>L728:CA<br>E730:CA | 1-25 | 26-50 | 51-75 |

| CAT-BRI1 interaction **(17 distances)** | | | | | |
|---|---|---|---|---|---|
| Y597:OH | CAT:H28 | 76 | Y597:OH | CAT:H4 | 77 |
| Y597:OH | CAT:O22 | 78 | Y597:OH | CAT:O23 | 79 |
| S647:O | CAT:H28 | 80 | S647:O | CAT:H4 | 81 |
| S647:O | CAT:O22 | 82 | S647:O | CAT:O23 | 83 |
| S647:N | CAT:H4 | 84 | S647:N | CAT:O23 | 85 |
| K601:NZ | CAT:H5 | 86 | K601:NZ | CAT:O6 | 87 |
| N705:OD1 | CAT:H28 | 88 | N705:OD1 | CAT:O3 | 89 |
| Y642:OH | CAT:H7 | 90 | R640:NH1 | CAT:O2 | 91 |
| Y599:OH | CAT:O6 | 92 | | | |
